# Characterization of methylation profiles in spontaneous preterm birth placental villous tissue

**DOI:** 10.1101/2021.04.26.441471

**Authors:** Heather M Brockway, Samantha L Wilson, Suhas G Kallapur, Catalin S Buhimschi, Louis J Muglia, Helen N Jones

## Abstract

Preterm birth is a global public health crisis which results in significant neonatal and maternal mortality. Yet little is known regarding the molecular mechanisms of idiopathic spontaneous preterm birth, and we have few diagnostic markers for adequate assessment of placental development and function. Previous studies of placental pathology and our transcriptomics studies suggest a role for placental maturity in idiopathic spontaneous preterm birth. It is known that placental DNA methylation changes over gestation. We hypothesized that if placental hypermaturity is present in our samples, we would observe a unique idiopathic spontaneous preterm birth DNA methylation profile potentially driving the gene expression differences we previously identified in our placental samples. Our results indicate the idiopathic spontaneous preterm birth DNA methylation pattern mimics the term birth methylation pattern suggesting hypermaturity. Only seven significant differentially methylated regions fitting the idiopathic spontaneous preterm birth specific (relative to the controls) profile were identified, indicating unusually high similarity in DNA methylation between idiopathic spontaneous preterm birth and term birth samples. We identified an additional 1,718 significantly methylated regions in our gestational age matched controls were the idiopathic spontaneous preterm birth DNA methylation pattern mimics the term birth methylation pattern, again indicating a striking level of similarity between the idiopathic spontaneous preterm birth and term birth samples. Pathway analysis of these regions revealed differences in genes within the WNT and Cadherin signaling pathways, both of which are essential in placental development and maturation. Taken together, these data demonstrate that the idiopathic spontaneous preterm birth samples are molecularly more mature than expected given their respective gestational age which likely impacts birth timing.

## Introduction

Preterm birth (PTB), defined as delivery at less than 37 weeks of gestation is the leading cause of neonatal mortality worldwide. Prematurity affects an average of 10% of infants born in the United States with rates increasing and costs approximately $26.2 billion dollars a year (annual societal cost including medical, educational, and lost productivity)(1, 2). The majority (50%) of preterm births are idiopathic and spontaneous (isPTB), rather than being medically indicated (e.g., pre-eclampsia). Risk factors include but are not limited to genetic ancestry, fetal sex, environmental exposures, and economic disparities(3). Complications include developmental delays, growth restriction, chronic respiratory problems as well as adult sequalae(3). Studies into the etiology of preterm birth have implicated a role for the placenta, a central component of the maternal-fetal interface, which has a vital role in pregnancy initiation, maintenance, and birth timing as well as fetal growth and development(4). As such, proper placental development, maturation, and function are essential for a successful pregnancy outcome and life-time offspring health. Each of these processes is an intricate balance of molecular interactions that are not fully understood even in healthy, normal, term pregnancies.

Placental maturation is accompanied by a marked increase in placental surface area due to placental remodeling initiated between 20-24 weeks gestation and continuing throughout the remainder of gestation which accommodates exponential fetal growth across the second half of gestation (4). Under normal physiological conditions, placental maturation is recognized by specific histological hallmarks including increased quantities of terminal villi (<80 microns in diameter), syncytial nuclear aggregates (SNAs, 10+ syncytial nuclei being extruded from the syncytiotrophoblast), and formation of the vasculosyncytial membranes (VSM) which when observed in significant quantities prior to 37 weeks, signify placentas with advanced villous maturation (AVM)(5, 6). Histological studies of pathological placentas indicate AVM occurs in 50-60% of isPTB and medically indicated preterm births(7, 8). This indicates a potential developmental disconnect between placental maturation and the corresponding fetal maturation. In infection associated preterm births, AVM was observed in less than 20% of pathologic placentas(7, 8). These studies indicate multiple morphological endotypes exist, underlying the classical clinical PTB phenotypes, especially those of spontaneous PTB which are based on gestational age and simply defined as early, moderate, and late(9). The identification of these morphological endotypes further highlights the heterogeneity confounding the identification of PTB etiology and potential diagnostic biomarkers.

Multiple levels of heterogeneity confound elucidation of molecular mechanisms involved in PTB, from inconsistent sampling of interface tissues to the numerous cell types within those tissues to individual differences within larger populations(10–13). However, traditional epidemiological studies have not accounted for this morphological, molecular, and physiological heterogeneity. Instead, the use of extensive covariate data to attempt overcome population-based heterogeneity has resulted in statistical overfit of models to their specific datasets, resulting in a loss of reproducibility and generalizability of biological inference across datasets(14, 15). This has led to a dearth of robust biomarkers capable of assessing spontaneous PTB risk and managing real-time clinical care. Our approach differs from the population based epidemiological approaches in that we focus molecular profiling in smaller, prescreened datasets with combined with select harmonizable covariate data that can be obtained for any dataset.

We have previously identified transcriptomic profiles of AVM in a small cohort using clinically phenotyped placental villous samples from spontaneous PTB births, including isPTB and infection associated births, between 29 and 36 weeks and normal term births (TB) between 38 and 42 weeks(16). In our datasets, we define infection associated preterm births as acute histologic chorioamnionitis (AHC) which have been identified via histological assessment of inflamed fetal membranes or molecular assessment(16). Given the importance of DNA methylation (DNAm) to placental development and maturation(17–20), we hypothesized the gene expression differences we observed in our transcriptome data could be due to changes in DNAm at CpG islands between the birth types. Therefore, we sought to identify specific DNAm profiles of placental maturation associated with our transcriptional profiles of maturation.

## Materials and Methods

### Study Population

This study was approved by the Cincinnati Children’s Hospital Medical Center institutional review board (#IRB 2013-2243, 2015-8030, 2016-2033). De-identified TB (n=6), isPTB (n=8), and AHC (n=8) placental villous samples along with appropriate covariate information were obtained from the following sources: The Global Alliance to Prevent Prematurity and Stillbirth (GAPPS) in Seattle Washington USA, the Research Centre for Women’s and Infant’s Health (RCWIH) at Mt Sinai Hospital Toronto Canada, and the University of Cincinnati Medical Center (UCMC). Inclusion criteria included: maternal age 18 years or older, singleton pregnancies with either normal term delivery (38-42 weeks’ gestation) or preterm delivery (29-36 weeks’ gestation) without additional complications. Additional information regarding these samples can be found in(16).

### Statistical Analyses

Cohort data were analyzed in Prism v8 (GraphPad). Data were evaluated for normality and non-parametric tests applied as appropriate. Non-parametric data are expressed as median and range and were analyzed by Kruskal-Wallis Test ANOVA with Dunn’s Multiple Comparisons. Categorical data were analyzed using Fisher’s Exact Test. These analyses were run independently of those included in(16).

### Intersection of transcriptomic candidate genes and CpG islands

Using the table function of the UCSC Genome Browser build hg38, we conducted a batch query using the 340 candidate genes from our previous transcriptome study(16). Using these genes as identifiers, we created an intersection with the CpG Island Track(21). This created an output table with gene names, genomic positions, and overlapping CpG islands. We then calculated the percentage of genes that overlapped with CpG islands.

### DNA Methylome Generation

DNA was isolated from homogenized, snap frozen placental villous samples using the DNAeasy Kit (Qiagen). DNA quantity and quality was assessed using Qubit 4 Fluorometer (Invitrogen) and Nanodrop Spectrophotometer (Thermo Fisher Scientific). A minimum of 500ng was submitted to the University of Minnesota Genomics Center and the University of Cincinnati Genomics, Epigenomics and Sequencing Core where DNA quantity and quality assessment on a Bioanalyzer (Aligent), bisulfite conversion, and methylome generation on the Illumina Methylation EPIC Bead Chip.

### DNA Methylation array data processing

Methylation data processing and analyses based on a previously developed workflow(22). All packages are available within Bioconductor(23) and all package scripts were run in RStudio/R v4.0.2(24, 25). IDAT file preprocessing and probe quality control was conducted in R using scripts based on minfi(26) and methylumi(27). IDAT files and a sample file containing covariate and BeadChip metadata were loaded into R where data quality was assessed using the mean detection p-values for the probes in each sample. We applied Functional Normalization(preprocessFunnorm)(28) for the algorithm’s ability to utilize the internal control probes for each individual sample in an unsupervised manner to control for unwanted variation.

After normalization, we excluded individual low-quality probes with a detection p-value > 0.1 in more than 2 samples or bead count <3 in at least 5% of samples, sex chromosome probes, cross-hybridizing probes, and probes where SNPs (within the binding region or within 5-10bp of the binding region) could potentially affect hybridization(22). To ensure appropriate filtering of problematic probes, we utilized several resources including the Illumina Methylation EPIC BeadChip hg38 manifest and Zhou et al(29) to identify additional variation that would interfere with probe hybridization. We utilized McCartney et al(30) to filter the cross-hybridizing probes that are not listed in the manifest. We removed all probes that reside in the ENCODE DAC black-list regions(31). All filtering criteria and number of probes filtered can be found in S1 Table.

Once probe filtering was complete, we assessed the data for batch effects using principal component analysis (PCA) and no significant batch effect was observed, therefore no correction was applied(32). The resulting data matrix contained M-values which were utilized for the statistical analyses of the pairwise comparisons due to their statistical robustness. β-values, which are transformed M-values, represent the ratio of all methylated probe intensities over total signal intensities or a percentage of methylation(33). All methylation values are delta M-values unless otherwise stipulated as they provide a better detection and true positive rates while reducing heteroscedasticity for methylation sites that are highly or non-methylated(33)

### Identification of differentially methylated positions

To assess differentially methylated positions (DMPs), we utilized generalized linear models within limma(34) to assess differential methylation for each individual probe within the M-value matrix as in (22) with adjustment for birth types and fetal sex as covariates within model. Due to the small sample numbers in our dataset, we did not assess any additional covariate data in this analysis as to not overfit the statistical models to this specific dataset and to increase generalizability of our findings in future studies. The following pairwise comparisons were used to identify significant positions of differential methylation: isPTB versus AHC, TB versus AHC and isPTB versus TB. The resulting output for these comparisons is a delta M-value representing the statistical difference in methylation at that position between the conditions being compared. Multiple corrections testing was conducted using the Benjamini Hochberg method(35) at multiple Q values: <0.05, <0.1, <0.2 and <0.3 (S2 Table). We tested Q values to determine if our lack observations in one pairwise comparison at Q=0.05 were due to a technical error or if these represented a true lack biological variability despite the statistical parameter selection. We opted to define significant DMPs with a Q <0.3 and a log2 fold-change of >±1.

### Methylome profile Identification

To identify methylation profiles, we used Venny 2.0(36) to generate Venn diagrams to intersect significant DMPs from each pairwise comparison to identify profiles specific to isPTB and AHC. An isPTB profile was defined as any DMP where the delta M-value of isPTB vs TB or AHC was differentially methylated compared to the delta M-values of AHC vs TB which were non-significant. An AHC profile was defined as any DMP where the AHC vs TB or AHC delta M-value was differentially methylated from the isPTB vs TB delta M-values which were non-significant. Heatmaps were generated in Prism v8 (GraphPad) using delta M-values. To assess if the differential methylation was influenced by outliers or by artifacts, we generated violin plots with β-values with median and quartiles in Prism v8 to check the distribution within selected individual samples.

### Differentially Methylation Region (DMR) Identification

We used DMRcate v2.2.3(22, 37) to identify differentially methylated regions comprised of significant DMPs within a specified distance using moderated t statistics. To identify significant DMPs within DMRcate, we used the M-value matrix (normalized and filtered) and set a threshold of Benjamini Hochberg adjusted p-value <0.3. Since DMRcate uses limma to determine the significant DMPs, we were able to utilize the same glm design from the initial DMPs analysis against adjusting for fetal sex. Once significant DMPs were identified, DMR identification thresholds were set at lamba=1000, C=2, and minimum cpgs=5. As we are analyzing array data, we opted to use the default lambda and C (scaling factor) which allows for optimal differentiation with 1 standard deviation of support to account for Type 1 errors. Once significant DMRs were identified in each pairwise comparison, we intersected them using Venny 2.0 to identify isPTB and AHC specific DMRs. The isPTB profile was defined as any DMR that was differentially methylated when compared to the AHC and TB, with the AHC vs TB. The AHC profile was defined as any DMR that was differentially methylated compared to isPTB and TB and where the isPTB vs TB methylation was non-significant meaning no DMR was identified in DMRcate. We also set a mean difference in differentiation threshold of 0.01. Heatmaps were generated in Prism v8 (GraphPad) using delta M-values.

### Functional analyses of DMRs with associated genes

Genes with associated DMRs were entered into the Panther Pathway DB(38) for statistical overrepresentation analyses for Reactome Pathways and to assess gene ontology (GO) for biological and molecular processes. Fisher’s Exact tests were used to determine significance and Bonferroni correction for multiple comparisons. Pathways were considered significant if they had an adjusted p-value <0.05.

### Intersection of DMRs with transcriptome candidate genes

To determine if any of our significant DMR’s impacted candidate gene expression, we intersected the DMR’s genomic locations with our candidate gene locations. All genomics regions were mapped to hg38. Where there was overlap, indicating a potential regulatory event, we took those locations and intersected with using the UCSC Genome Table browser (hg38) and the CpG island tracks (21), using the feature-by-feature function. This allowed for identification of DMRs in CpG regions of our candidate genes.

## Results

### Methylation Study Characteristics

Maternal and fetal characteristics for the three different pregnancy outcomes included in the DNAm analyses are presented in Table 1. Transcriptomes from these samples were previously published(16). Due to the amount of sample required for DNA extraction only a subset of the samples were used and the statistical analyses repeated but did not change. Significant differences were observed in gestational age and fetal weights between AHC and isPTB samples compared to the TB samples (p<0.05). All AHC and TB for which there were fetal weights available were appropriate for gestational age. We included males and females in each sample set and adjusted the linear models for fetal sex in addition to birth outcome. It is important to note that in this study, we have mixed genetic ancestry within each of the sample sets.

**Table 1:**
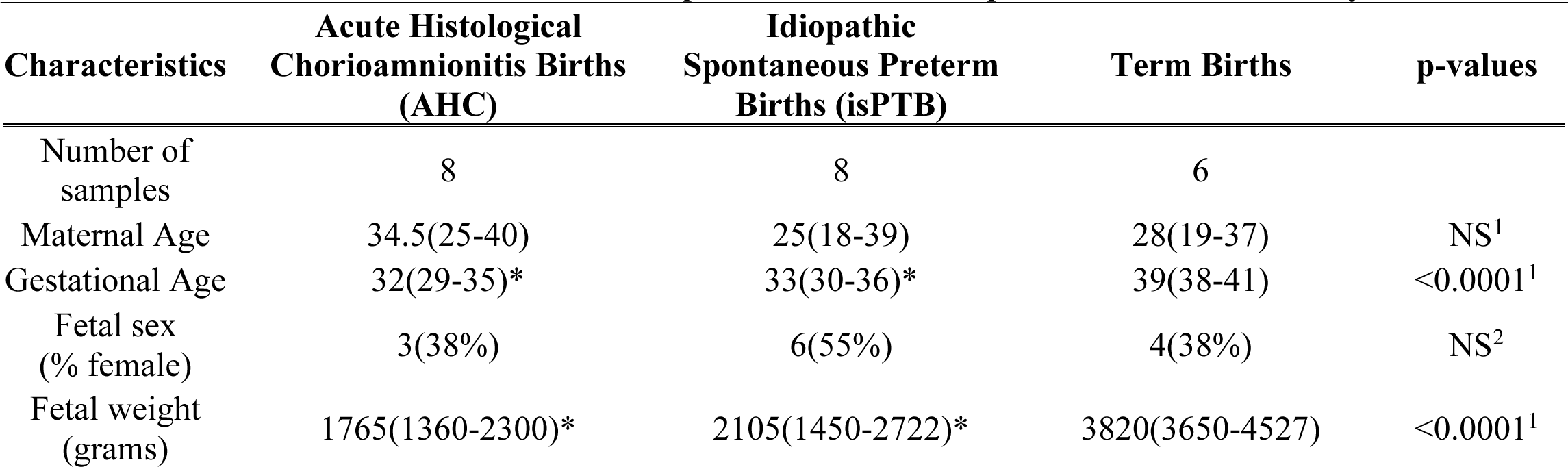
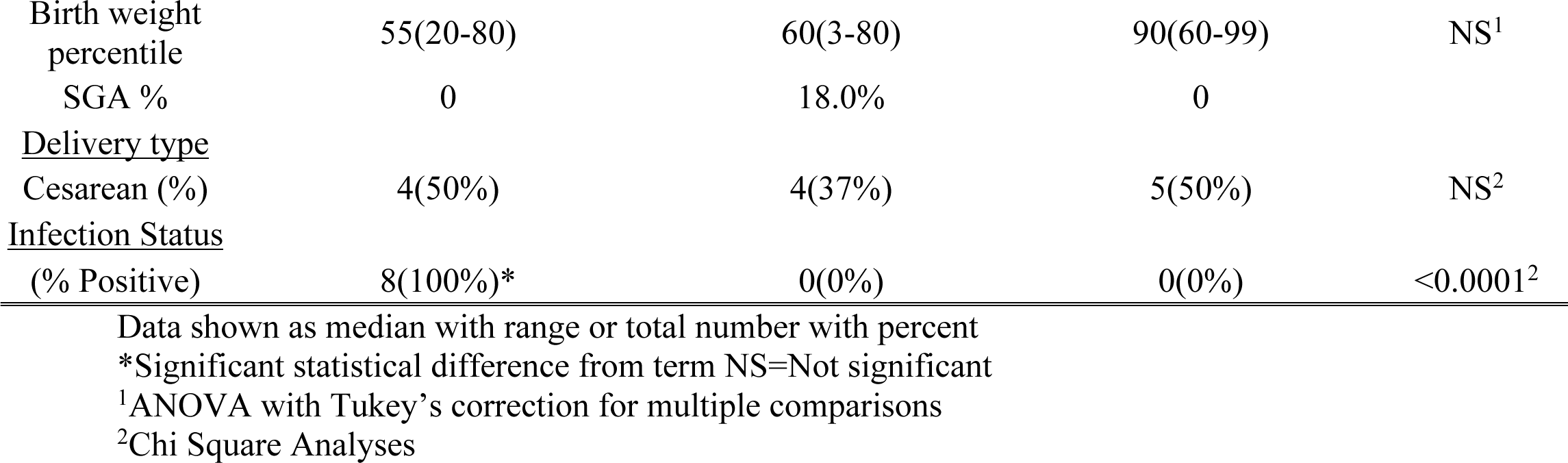
Clinical characteristics of the placental villous samples included in this study

### Identification of transcriptomic profile candidate genes with overlapping CpG islands

The intersection of isPTB specific methylation profiles with the previously identified 170 upregulated genes in isPTB samples yielded 102 candidates (60%) overlapping with CpG islands in their coding regions. In the AHC profile, 120/170 (81%) candidate genes intersected with CpG islands within coding regions.

### Identification of significant differentially methylated positions (DMP)

Preliminary quality control identified one sample with mean probe detection p-value >0.1 and it was subsequently removed from methylation analyses. Prior to normalization and subsequent probe filtering, there were 866,901 probes in the data matrix. After normalization and filtering, 108,691 probes were removed, leaving 758,210 probes in the matrix for analyses (S1 Table).

Our initial statistical testing using the Benjamini Hochberg Q cutoff of 0.05 did not yield any significant DMPs in the isPTB vs TB pairwise comparison. With a Type 1 error rate of 5%, we expected to observe approximately 37,910 statistically significant DMPs in this comparison; however, we observed 0. By relaxing the rate of acceptable Type 1 errors to 30%, we would expect to observe 227, 463 statistically significant DMPs, yet we only observed a total of 662 significant DMPs (S2 Table). We test modeled various statistical parameters to determine if our observations were due to technical errors or true biological differences. At every Q value tested and with different statistical models, we observed the number of DMPs between isPTB and TB to be significantly less than expected. Ultimately, we opted on a Q cutoff of 0.3 in limma(34).

We then set a threshold for differential methylation of log2 fold-change of >1. The DMP analysis identified a total of 24,202 significant DMPs across all pairwise comparisons in the model. In the isPTB vs AHC comparison we identified 8,309 DMPs, 4,334 with reduced methylation and 3,975 more methylated in isPTB compared to AHC. In the TB vs AHC comparison, we identified a total of 15,817 DMPs with 7,170 less methylated and 8,647 more methylated in TB. Lastly, in the isPTB vs TB comparison, 85 DMPs were identified as significant with 13 more methylated and 72 less methylated (Fig 1A).

**Fig 1:**
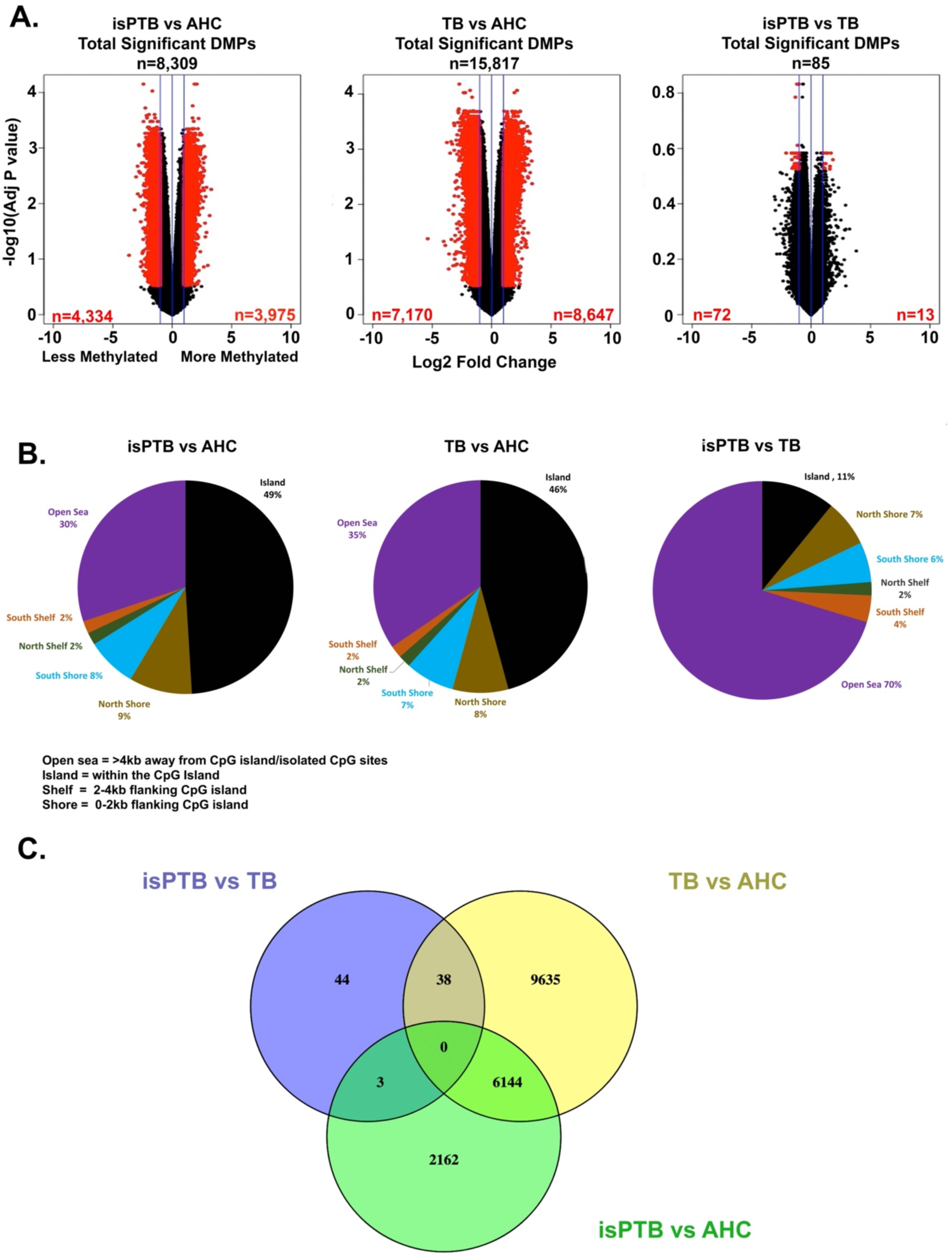
Identification of methylation profiles using a comparative approach. **A.** Differentially methylated positions were identified using pairwise comparisons in limma. Red points indicate significant DMPs with a threshold of log2 fold-change >1 and Benjamini Hochberg adjusted p-value <0.3. Blue lines represent log2 fold-change of 1. **B.** Genomic distribution of DMPs in the pairwise comparisons. The majority of DMPs in the isPTB and TB verses AHC comparisons are located inside or close to known CpG islands. However, in the isPTB verses TB comparison, the majority of DMPs are in open sea regions with no known islands within 4kb. **C.** The venn diagram represents the intersection of pairwise comparisons to classify significant DMPs into isPTB and AHC specific methylation profiles.

We observed differences in genomic location of the DMPs between the pairwise comparisons and thus, analyzed the genomic location distribution of the DMPs per comparison (Fig 1B). In the isPTB vs AHC and TB vs AHC comparisons the majority of DMPs were associated with CpG islands, shores, shelves (isPTB = 70% and TB = 65%) while the remaining DMPs were in open sea locations which are typically 3-4kb away from CpG islands (isPTB = 30% and TB = 35% respectively). In contrast, in the isPTB vs TB comparison, 70% of the DMPs were associated with open sea positions while only 30% associated with CpG islands, shores, and shelves. The first step in identification of a DMP methylation profile was to intersect the significant DMPs from each pairwise comparison and determine which would potentially segregate into an isPTB or AHC profile (Fig1C).

### Isolation of isPTB and AHC DNA methylation profiles using DMPs

As a result of the intersection of significant DMPs, we identified 47 potential isPTB specific DMPs. Upon examining the DNAm patterns for these DMPs across all pairwise comparisons, we wanted to know which DMPs has differential methylation in the isPTB versus the AHC and TB. We ultimately isolated 3 isPTB specific DMPs out of the 47 potential isPTB DMPs. Our examination of the individual sample beta values and their distribution for each DMP confirmed our findings were not due to artifacts or outliers (Fig 2A). Although we initially identified 8,306 potential AHC specific DMPs via the intersection, upon further examination of the DNAm pattern, we ultimately isolated 6,177 where the AHC samples were differentially methylated compared TB or isPTB (Fig 2B). Of these, 3,002 are more methylated and 3,175 are less methylated. We also examined the genomic location distribution of the AHC profile DMPs and found that 76% were located within CpG islands, shores, and shelves with remaining 24% located in open sea regions (S1 Fig).

**Figure 2:**
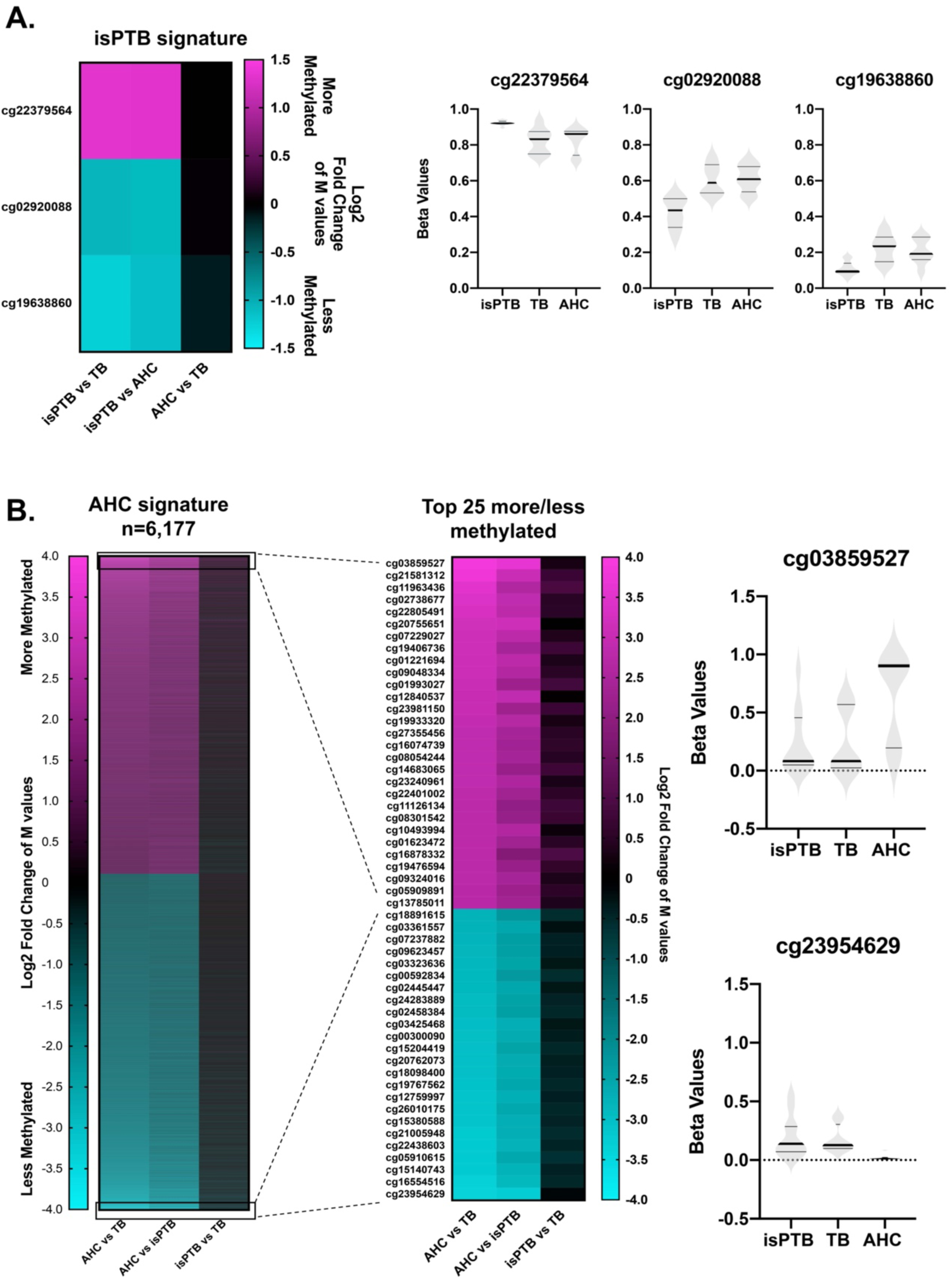
Identification of significant methylation profiles for isPTB and AHC DMPs. **A.** Three DMPs identified as having an isPTB specific methylation pattern where the isPTB samples were differentially methylated compared to the AHC or TB samples. The distribution of individual sample beta values was assessed to determine if there were outliers or artifacts influencing the methylation patterns. The dark bands represent the mean of the methylation values while the lighter grey bands represent the interquartile range. **B.** 6,177 DMPs demonstrating a methylation pattern where the AHC samples were differentially methylated compared to the isPTB or TB samples. The breakout heatmap shows the pattern or the top 25 more and less methylated samples and demonstrates the similarity of methylation between the isPTB and TB samples. The distribution of individual sample beta values was assessed to determine if there were outliers or artifacts influencing the methylation patterns.

### Identification of differentially methylated regions (DMRs)

To identify differentially methylated regions, we used the M-value matrix of data values previously generated in our initial analyses. We utilized again a relaxed Q <0.3 to ensure we would be able to identify enough CpG sites to identify DMRs in the isPTB vs TB comparison (S3 Table). Only then, we were able to identify significant DMRs within all pairwise comparisons (Table 2).

**Table 2.** Summary of significantly differentiated DMRs identified by DMRcate encompassing both coding and non-coding loci

| Pairwise comparison | Number of Significant DMRs Identified* | Width of DMR (Range) | Number of Significant Probes in DMR (Range) |
| --- | --- | --- | --- |
| isPTB vs TB | 56 | 180-1750bp | 5-18 probes |
| isPTB vs AHC | 12,883 | 83-9,386bp | 5-110 probes |
| TB vs AHC | 19,006 | 37-14,383bp | 5-202 probes |

\*minimum smoothed FDR <0.05

56 DMRs were observed within the isPTB vs TB comparison in contrast to the thousands significant DMRs identified in the isPTB and TB verses AHC pairwise comparisons. All isPTB vs TB DMRs were under 2000bp wide and had no more than 18 CpG sites in any given DMR. In contrast, the DMRs in the isPTB and TB vs AHC comparisons were wider and encompassed more probes (Table 2). We intersected the DMRs and identified potential candidate DMRs for isPTB and AHC methylation profiles (S2 Fig). Ultimately, we identified 51 potential isPTB specific and 12,843 AHC specific DMRs. These DMRs overlap with coding and non-coding loci across the genome as per the annotation from DMRcate package(37).

### Identification and function of DMRs specific to isPTB and AHC

Of the 51 candidate isPTB DMRs, only seven demonstrated an isPTB specific profile (Fig 3 and Table 3). Six isPTB specific DMRs overlap coding/non-coding loci with only one sitting in an upstream promoter region, *LINC02028* (Table 4). This is the only isPTB-specific DMR that overlaps with a CpG island. Four of the DMRs sit within transcripts for *FAM186A, NOD2, UBL7-AS1*, and *PDE9A*, more specifically within introns or at intron/exon boundaries. The remaining two DMRs sit in the 3’UTR of genes, *ZBTB4 and STXB6*, with the *ZBTB4* DMR crossing the last exon/UTR boundary (Table 4). No over-represented pathways were identified.

**Figure 3:**
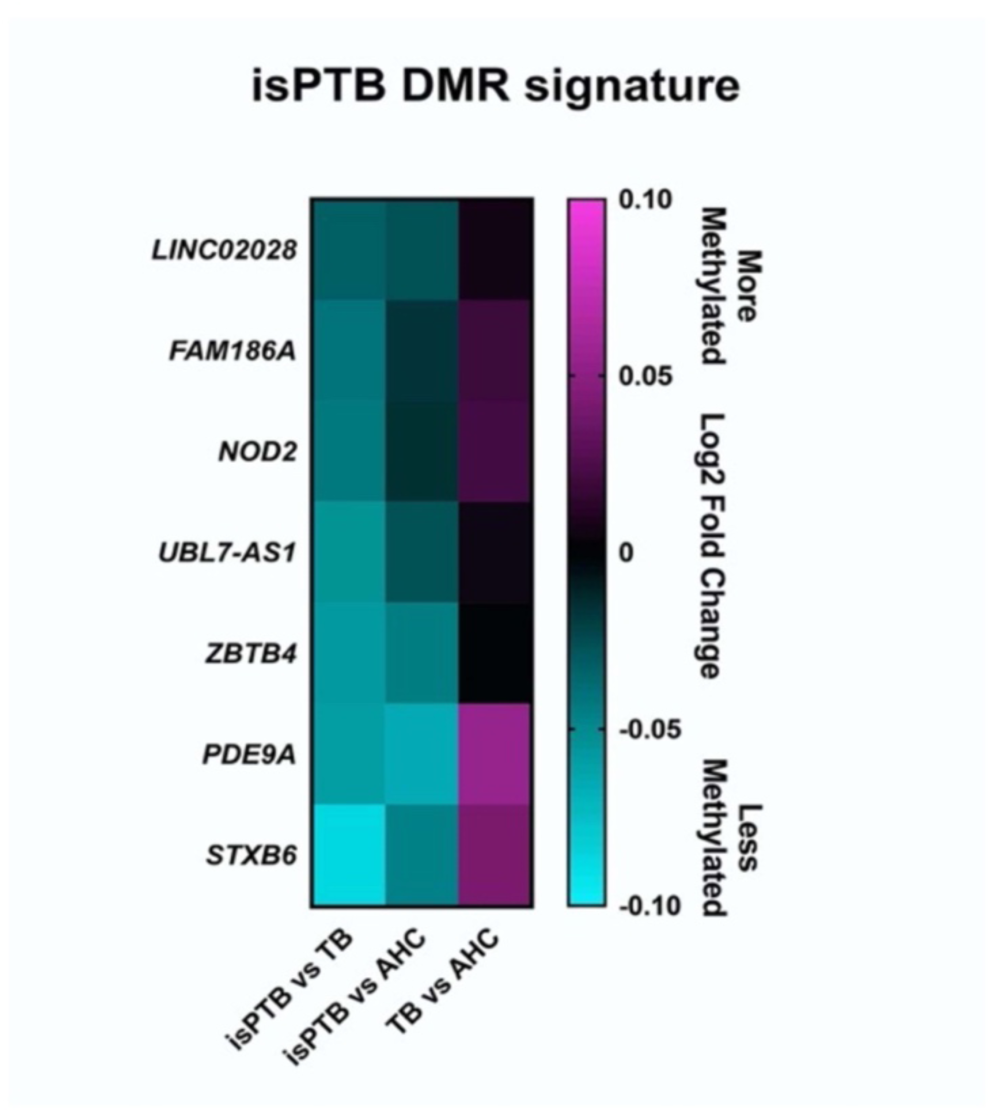
isPTB specific DMR profile. Differentially methylated DMRs were identified by differences in the mean of the probe values across the DMR. Only 7 isPTB DMRs had an isPTB specific profile where the isPTB DMRs were less methylated than the TB or AHC DMRs. Two of the DMRs overlap non-coding regions. No DMRs were identified that were more methylated.

**Table 3:** Summary of isPTB profile DMRs

| Locus | Mean Difference Methylation for all probes in DMR |  |  | DMR coordinates |
| --- | --- | --- | --- | --- |
|  | isPTB vs TB | isPTB vs AHC | TB vs AHC |  |
| *LINC02028* | -0.033 | -0.028 | 0.007 | chr3:194072066-194072416 |
| *FAM186A* | -0.0416 | -0.0175 | 0.0192 | chr12:50343856-50344626 |
| *NOD2* | -0.043 | -0.015 | 0.022 | chr16:50715192-50715700 |
| *UBL7-AS1* | -0.054 | -0.028 | 0.005 | chr15:74466794-74467158 |
| *ZBTB4* | -0.058 | -0.045 | 0.0008 | chr17:7461421-7462028 |
| *PDE9A* | -0.059 | -0.066 | 0.054 | chr21:42733397-42733894 |
| *STXB6* | -0.087 | -0.0466 | 0.042 | chr14:24808650-24810213 |

**Table 4:** Functional information for the isPTB DMRs

| Locus | Overlaps with CpG Island | Location |
| --- | --- | --- |
| *LINC02028* | chr3:194070715-194071468 | Promoter |
| *FAM186A* | NA | Intronic |
| *NOD2* | NA | Intronic |
| *UBL7-AS1* | NA | Intronic |
| *ZBTB4* | NA | 3'UTR/last exon |
| *PDE9A* | NA | Intron/exon boundary |
| *STXB6* | NA | 3'UTR |

Of the 12,843 AHC specific DMRs, only 1,718 demonstrated an AHC specific methylation pattern. These DMRs include coding and non-coding loci (Fig 4A and S4 Table). Of these, 801 DMRs are more methylated while 917 are less methylated than corresponding DMRs in the isPTB or TB pairwise comparison. In the top 25 more/less methylated loci, the lack of significant differences in methylation can clearly be observed in TB vs isPTB (Fig 4B). Of these, 19% (n=328) had direct overlap with CpG islands. The remaining 81% had no overlap at all with CpG islands.

**Figure 4:**
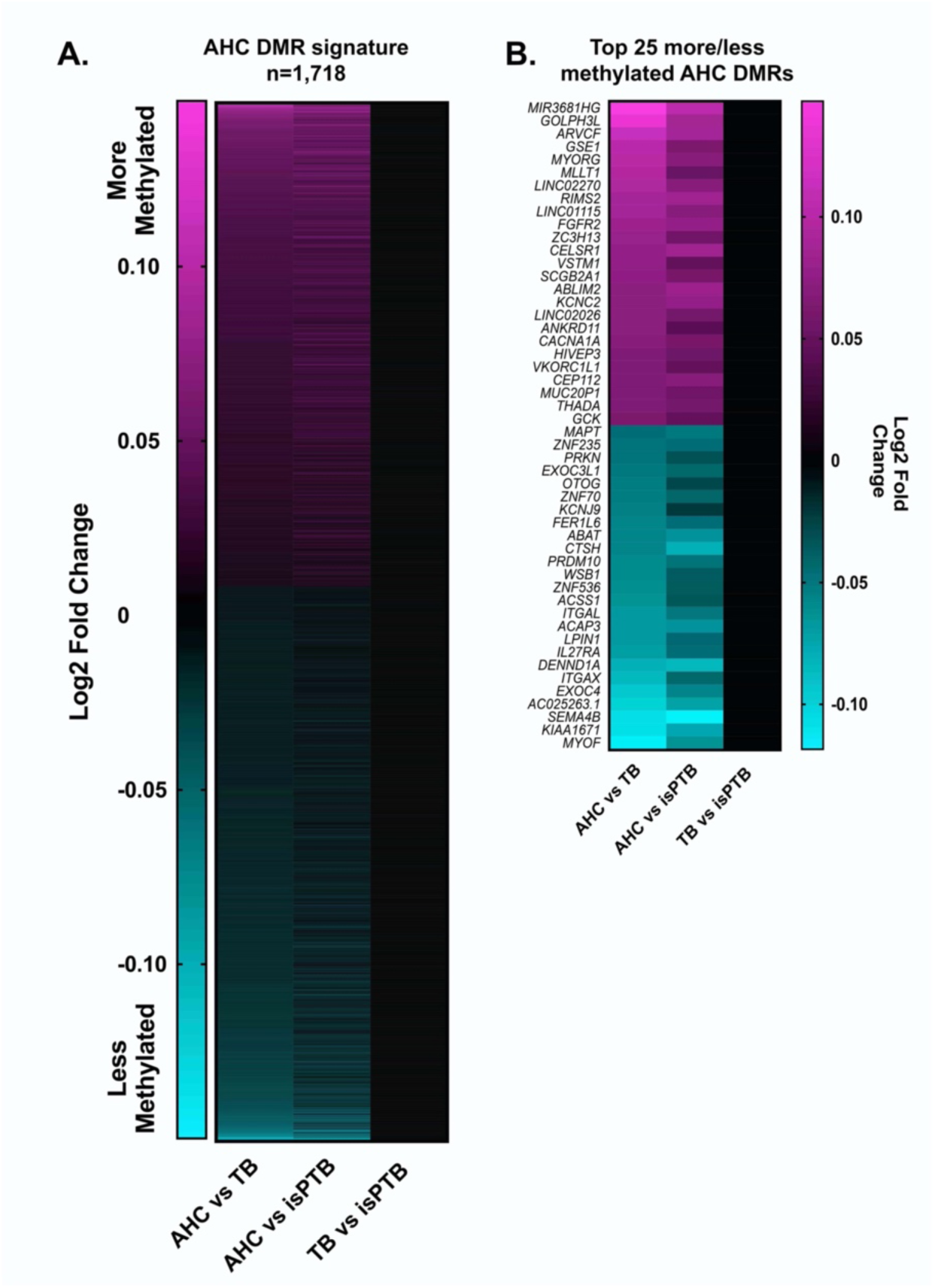
AHC specific DMR profile. **A.** Differentially methylated DMRs were identified by differences in the mean of the probe values across the DMR. AHC specific DMRs are defined by when the AHC DMRs were differentially methylated compared to the TB or isPTB DMRs. **B.** The top 25 more and less methylated DMRs demonstrates the clarity of the molecular profile, as there is no significant differential methylation in the TB vs isPTB comparison.

We assessed the potential implications of the AHC specific DMRs using statistical over-representation analyses for pathways and GO terms. In the more methylated DMRs, we identified two significantly over-represented pathways: WNT and Cadherin signaling (Table 5). Significant Biological Process GO terms included homophilic cell adhesion via plasma membrane adhesion molecules (GO:0007156) and cell-cell adhesion via plasma-membrane adhesion molecules (GO:0098742).

**Table 5:** Bioinformatic functional assessment of more methylated AHC profile DMRs via PantherDB

|  | *Homo sapiens*<br>(all genes in<br>database) | Genes from<br>input list | Expected | Fold<br>Enrichment | Adjusted<br>p-value* |
| --- | --- | --- | --- | --- | --- |
| **PANTHER Pathways** |  |  |  |  |  |
| Cadherin signaling<br>pathway (P00012) | 164 | 21 | 5.34 | 3.94 | 6.51E-05 |
| Wnt signaling pathway<br>(P00057) | 317 | 30 | 10.31 | 2.91 | 1.03E-04 |
| **GO biological process**<br>**complete** |  |  |  |  |  |
| homophilic cell adhesion<br>via plasma membrane<br>adhesion molecules<br>(GO:0007156) | 168 | 26 | 5.47 | 4.76 | 4.62E-06 |
| cell-cell adhesion via<br>plasma-membrane<br>adhesion molecules<br>(GO:0098742) | 257 | 28 | 8.36 | 3.35 | 1.05E-03 |
| **GO molecular function**<br>**complete** |  |  |  |  |  |
| ion binding<br>(GO:0043167) | 6354 | 277 | 206.71 | 1.34 | 5.61E-05 |
| binding (GO:0005488) | 16539 | 593 | 538.05 | 1.1 | 8.90E-05 |
| molecular function<br>(GO:0003674) | 18245 | 631 | 593.55 | 1.06 | 4.23E-03 |
| metal ion binding<br>(GO:0046872) | 4268 | 192 | 138.85 | 1.38 | 4.82E-03 |
| cation binding<br>(GO:0043169) | 4354 | 194 | 141.65 | 1.37 | 9.08E-03 |
| adenyl nucleotide binding<br>(GO:0030554) | 1572 | 84 | 51.14 | 1.64 | 3.90E-02 |

\*Fisher Test Bonferroni Corrected for multiple comparisons

No significant over-represented pathways were identified in the less methylated DMRs. The significant Biological Process GO terms that were associated with the less methylated dataset include cell morphogenesis involved in differentiation (GO:0000904), cell morphogenesis (GO:0000902) and detection of chemical stimulus (GO:0009593). For Molecular Function, the following significant GO terms were identified: ion binding (GO:0043167), protein binding (GO:0005515), protein binding (GO:0005515), and olfactory receptor activity (GO:0004984) (Table 6)

**Table 6:**
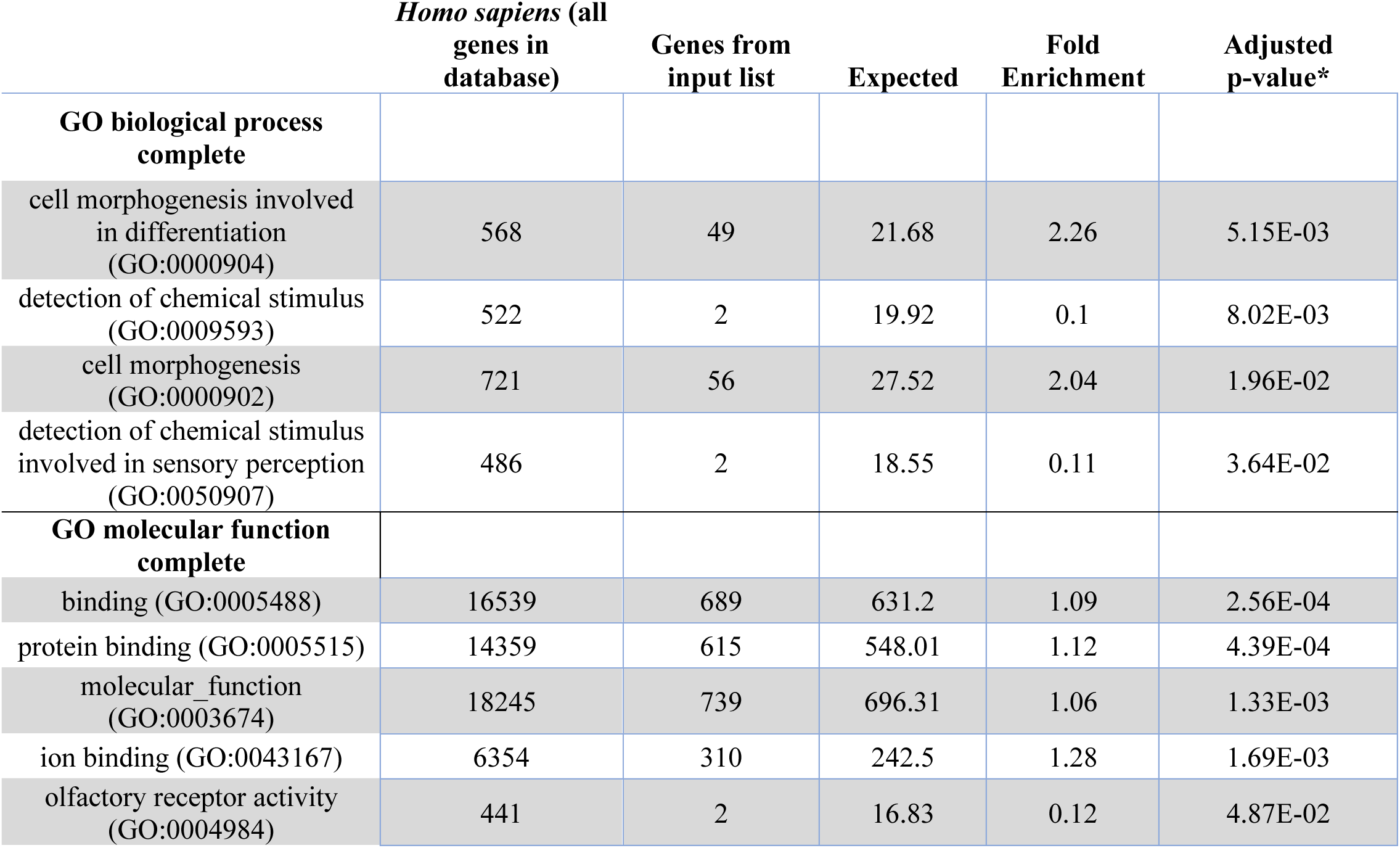
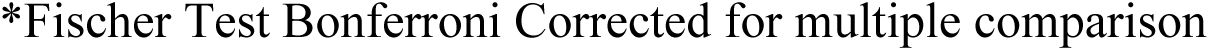
Bioinformatic functional assessment of less methylated AHC profile DMRs via PantherDB

|  | *Homo sapiens* (all<br>genes in<br>database) | Genes from<br>input list | Expected | Fold<br>Enrichment | Adjusted<br>p-value* |
| --- | --- | --- | --- | --- | --- |
| **GO biological process**<br>**complete** |  |  |  |  |  |
| cell morphogenesis involved<br>in differentiation<br>(GO:0000904) | 568 | 49 | 21.68 | 2.26 | 5.15E-03 |
| detection of chemical stimulus<br>(GO:0009593) | 522 | 2 | 19.92 | 0.1 | 8.02E-03 |
| cell morphogenesis<br>(GO:0000902) | 721 | 56 | 27.52 | 2.04 | 1.96E-02 |
| detection of chemical stimulus<br>involved in sensory perception<br>(GO:0050907) | 486 | 2 | 18.55 | 0.11 | 3.64E-02 |
| **GO molecular function**<br>**complete** |  |  |  |  |  |
| binding (GO:0005488) | 16539 | 689 | 631.2 | 1.09 | 2.56E-04 |
| protein binding (GO:0005515) | 14359 | 615 | 548.01 | 1.12 | 4.39E-04 |
| molecular function<br>(GO:0003674) | 18245 | 739 | 696.31 | 1.06 | 1.33E-03 |
| ion binding (GO:0043167) | 6354 | 310 | 242.5 | 1.28 | 1.69E-03 |
| olfactory receptor activity<br>(GO:0004984) | 441 | 2 | 16.83 | 0.12 | 4.87E-02 |

\*Fischer Test Bonferroni Corrected for multiple comparison

### Identification of DMRs in regulatory elements of transcriptome candidate genes

Upon intersection of significant DMRs and the candidate genes, none of the isPTB DMRs intersected with any of the isPTB candidate genes. Out of the 1,718 significant AHC DMRs, only eight intersected with the AHC candidate genes (Table 7). Interestingly, six of these DMRs have methylation patterns, in all cases less methylated, that agree with upregulated transcription status. The remaining two have no correlation between profiles (S5 Table).

**Table 7:**
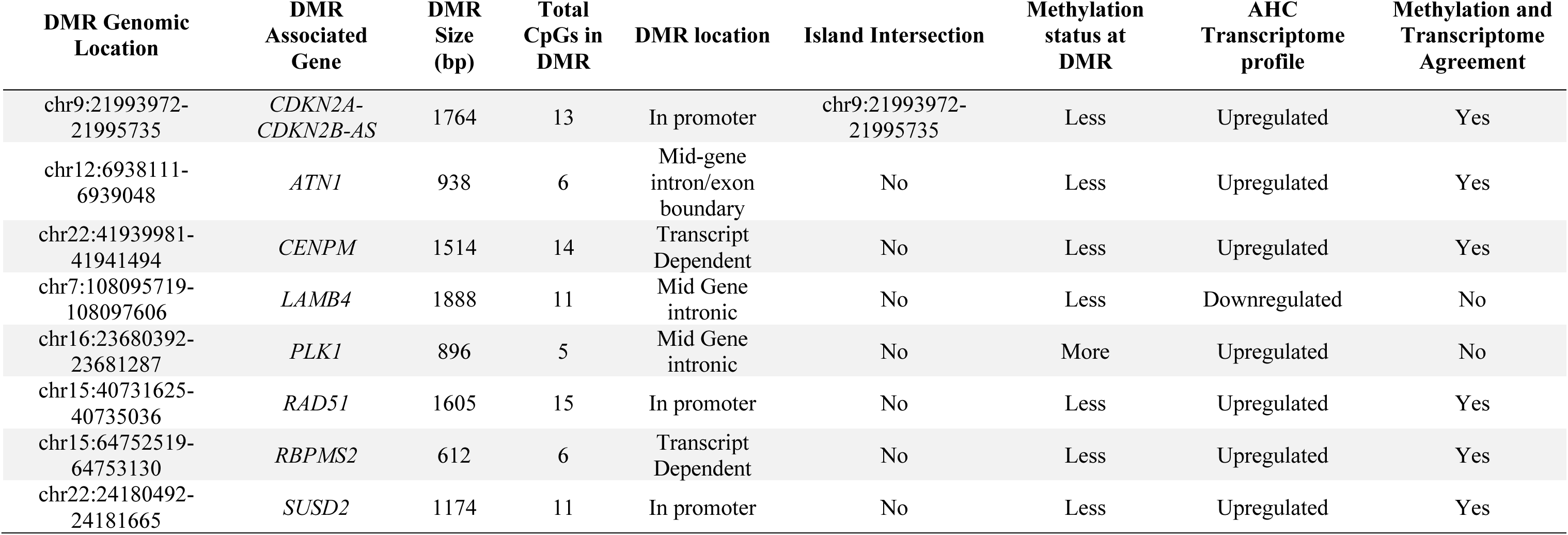
DMR Characterization and comparison to transcriptome profiles

For each of these eight genes, we examined the genomic location to determine if these DMRs were in promoters or CpG islands, potentially regulating gene expression. We observed only one DMR, *CDKN2A,* that overlapped with CpG islands 5’ upstream of their transcripts. The DMR upstream of *CDKN2A* also resides in the same genomic area as a non-coding transcript, *CDKN2B*. The remaining seven DMRs did not overlap any CpG islands although, two were in the promoter or first intronic region of their associated genes. *CENPM* and *RBPMS2* have multiple transcripts and the location of the DMR varies depending on the specific transcript length and start site. Three DMRs reside in introns or across intron/exon boundaries (Table 7).

## Discussion

To gain insight into the role of DNA methylation in spontaneous preterm birth, we utilized pairwise comparisons of methylation between spontaneous preterm births and normal term births using a general linear model adjusting for fetal sex and gestational age at delivery. It is essential to note that normal gestationally age-matched placental samples are typically not available for studies such as this depending on ethical restrictions of the geographical locale of the study. Therefore, we opted to use with acute histologic chorioamnionitis samples (AHC) which been previously shown to have much lower occurrences of AVM than other clinically defined preterm birth types including PE and IUGR(7, 8) We were able to identify distinct methylation profiles at both the positional (DMP) and regional (DMR) levels in isPTB and AHC. Through bioinformatic functional assessment, we were able to identify pathways of interest pertaining to placental maturation.

Our preliminary analyses indicated that there were very few DMP and DMR between the isPTB and TB birth types regardless of the statistical parameters applied. We tested multiple parameters within the statistical models to ensure that lack of differences was likely due to biological factors, not technical errors. Given the sheer number of datapoints being examined, we felt that relaxing the Q value to 0.3 would not adversely affect our analyses and we were willing to accept the potential increase in false positives(39, 40). This allowed us to better assess any potential differences between isPTB and TB despite the potential increase in false positives. The Benjamini Hochberg correction is dependent on the overall number of samples to be corrected and considered to be rather conservative. Regardless of the statistical parameters applied, the isPTB profile mimicked the TB profile to a high degree which, agrees with the transcriptomic profiles we previously identified(16) and provides additional evidence of a potential placental hypermaturity profile associated with isPTB. Although this a preliminary study investigating DNA methylation in spontaneous preterm birth, this pattern of DNA methylation was also observed in studies of iatrogenic preterm births in DMP and DMR analyses, for both PE and IUGR(20). In the second study, focusing on imprinted regions found that IUGR samples also mimicked the PE and term controls(41). Pyrosequencing from this second study confirmed no differences in the DMRs suggesting the detection of hypermaturity molecular profile. Given that hypermaturity is estimated to affect 50-60% of all preterm births including PE and IUGR(7, 8), these results provide additional evidence supporting the use of placental DNAm clinically to classify pathophysiologies such as hypermaturity(20, 42).

DMRs are associated with numerous disease pathologies in multiple tissues(43, 44). While DNAm has been studied in the other adverse pregnancy outcomes such as PE and IUGR, this study is the first to look specifically at isPTB. Our analysis resulted in the identification of seven DMRs with isPTB specific methylation patterns; two are associated with non-coding transcripts (*LINC02028* and *UBL7-AS*), five with genes (*ZBTB4, STXBP6, PDE9A, NOD2, and FAM186A*). Of these genes, four are of particular interest due to their potential function in or previous association with PTB.

*ZBTB4* is a placentally expressed gene coding for a transcription factor that binds methylated CpGs in a repressive manner, controls TP53 responses in cells, and inhibits cell growth and proliferation (45–47). TP53 was identified as a potential biological pathway of interest in our microarray meta-analysis of spontaneous PTB(48) and has been implicated in isPTB from a uterine perspective in mice(49). *STXBP6,* also known as *AMISYN,* binds SNARE complex proteins together(50). As SNARE complexes have been well described in synaptic vesicle formation and exocytosis(51) and regulation of membrane fusion dynamics(52, 53), the presence of this protein in the placenta suggests potential role in placental extracellular vesicle formation or the mediation of membrane fusion during cytotrophoblast differentiation(52, 54).

*PDE9A* functions in the hydrolysis of cAMP into monophosphates, modulating the bioavailability of cAMP and cGMP in cells(55). cAMP signaling is essential to cytotrophoblast differentiation into syncytiotrophoblast(56); therefore, alteration of PDE9A expression or function impacts cAMP bioavailability potentially altering this specific trophoblast differentiation pathway. In fact, PDE9A has been proposed as a potential first trimester maternal serum biomarker for Trisomy 21(57). Placentas from Trisomy 21 fetuses have multiple defects in cytotrophoblast differentiation, specifically cell fusion, resulting in what appears to be delayed villous maturation, indicating a key role for this gene in normal placental maturation(57–60).

*NOD2* has a role in activation of the innate inflammatory response and has been implicated in NFKB activation(61–63). NFKB activation is a central component of pro-inflammatory /labor pathways in both normal term and preterm pathophysiology(62,64,65). As a member of the NOD-like receptor family, NOD2 has been previously associated with recognition of pathogen associated molecular patterns (PAMPs) and damage associated molecular patterns (DAMPs) both of which have been associated with preterm labor and birth(62). The activation of pathways associated with PAMPs and DAMPs have previously been associated with sPTB and iatrogenic PTB(48,66–68). NOD2 has been studied primarily in the context of a proinflammatory factor in fetal membranes and myometrium; however, *NOD2* is expressed in first trimester and term placental tissues, specifically in syncytiotrophoblast and stromal cells(61, 69). Furthermore, NOD2 polymorphisms have been associated with preterm birth in several genetic studies examining innate immunity, preterm premature rupture of membranes (PPROM), and early onset PE and HELLP (Hemolysis, Elevated Liver enzymes and Low Platelets) syndromes(62,67,70,71).

Taken together, these isPTB DMRs and their associated genes suggest that altered DNA methylation maybe highly influential in isPTB; however, from these data alone, it cannot be determined if this is causative or the result of isPTB as the samples were obtained at delivery. Although we cannot sample placental tissues throughout gestation to determine cause or effect, using DNAm profiling on delivered placental tissues will provide key insights in the pathophysiological underpinnings of adverse pregnancy outcomes.

In contrast to the isPTB DNAm profile, our examination of the AHC samples compared to the isPTB and TB samples identified 1,718 DMRs. We observed within the top 25 more/less methylated DMRs, multiple DMRs were associated with genes of interest that were previously associated with adverse pregnancy outcomes including IUGR and PE. Several have also been associated gestational diabetes mellitus (GDM) which can also result in preterm birth. These genes of interest include: *MLLT1*(72), *FGFR2*(72), *CACNA1A*(73), *GCK*(74, 75), *FER1L6*(76), *CTSH*(77), and *ACAP3*(78). Additionally, *GSE1* (79), *VSTM1*(80), and *ACSS1*(79) are expressed in the placenta but have not yet been associated with an adverse pregnancy outcome. Our pathway analyses of the more methylated DMRs, yielded two pathways with statistical over-representation, WNT and Cadherin signaling. Both pathways are necessary for placental development and maturation(81–84) and a prior methylation study in PE also identified differential methylation (increased methylation) in WNT and cadherin signaling(85), which agrees with our findings. Given that over 50% of PE cases have hypermaturity along with the pathological hallmarks of PE, this may indicate a role for these pathways in placental maturation.

We initially hypothesized that changes in methylation at CpG islands could be driving the transcriptional differences we previously observed. However, when we intersected our significant DMRs with our candidate genes, we did not observe any overlap in the isPTB profiles and only eight examples of overlap in the AHC profiles. Of those eight DMR/gene combinations, only *CDKN2A/CDKN2B-AS* overlapped with a CpG island. *CDKN2A,* also known as p16, is a gene with multiple transcripts which have different first exons. Known as an important tumor suppressor, its primary role is regulating cell cycle progression through the regulation of TP53. Loss of function studies of *Cdkn2a* and *Tp53* in mice have demonstrated histopathological changes in placenta and upregulated senescence markers as well as mitotic inhibition(86). *CDKN2B-AS* is a functional RNA with regulatory roles via interaction with *PRC1* and *PRC1* which regulates the rest of the genes in this locus epigenetically(87). Additionally, *CDKN2B-AS*, also known as *ARNI*L, has been implicated in preterm birth

Interestingly, This DMR resides in locus consisting of *CDKN2A/CDKN2A-DT/CDKN2B-AS/CDKN2B*, a locus vital to cell cycle control and is dysregulated in many cancers. *CDKNA-DT* is a divergent transcript with no known function. However, *CDKN2B*, also known as *p15,* is another critical tumor suppressor which inhibits cyclin kinases CDK4 and CDK6(87). These data along with our methylation data suggest the correct expression of the *CDKN2A/CDKN2A-DT/CDKN2B-AS/CDKN2B* locus is critical to the structure, function, and potentially the rate of maturity of the placenta and normal healthy pregnancy.

*CENPM* and *SUSD2* all have roles in cell cycling and proliferation with mutations associated with cancers. In many cancers the loss of methylation is associated with cell proliferation and migration via metastasis. However, in the developing and maturing placenta these processes are essential for growth, function, and maturation(42,88,89). Less methylation at the DMRs associated with *RAD51, RBPMS2, ATN1* and the corresponding upregulation could be indicative of senescence given their respective roles in DNA repair, regulation of cell differentiation, and transcriptional repression. While the intersection of our matched transcriptional and methylation data did not necessarily support our original hypothesis of gene regulation via CpG islands in promoter regions, we were able to identify a potentially critical biological function, cell proliferation and an essential locus, *CDKN2A/CDKN2A-DT/CDKN2B-AS/CDKN2B*, for further study.

One of the caveats to studying placental villous omics of any nature is the lack of normal gestational age matched tissue due to limited accessibility throughout gestation. We previously utilized infection associated samples in our transcriptome analyses as our gestational age controls as their villi did not appear to be inflamed via pathological assessment. While we cannot rule out that changes at AHC loci may be due to infection, we did not observe pathways or GO terms associated with immunity or infection. Our data suggests that the overall AHC DNAm profile is reflective of appropriate villous maturation rather than an infection profile as was observed in our transcriptome data(16).

This is the first study to examine DNAm in spontaneous preterm birth in the context of placental maturity. The identification of hypermaturity profiles by both positional and regional differences in methylation highlights importance of DNAm to placental maturation and thus warrants further study.These differences could be due to altered trophoblast biology. These data when taken in the context of a potential epigenetic clock, suggests that perhaps epigenetic aging may have a role as it has in other fetal tissue and stem cells(90, 91). Future studies need to investigate the origin of the observed hypermaturity and its impact on the maternal-fetal interface and pregnancy outcomes.

## Supporting information

Supplemental Fiiles

## Acknowledgements

The authors would like to express their gratitude to the patients who donated their placentas for research. We would also like to thank Pietro Presicce, Paranthaman Senthamarai Kannan, and Manuel Alvarez (Kallapur lab), William E. Ackerman, Irina A. Buhimschi (Buhimschi lab), GAPPS, and RCWIH for assisting us in obtaining the placental samples and covariate data. Additionally, the authors thank the staff at the University of Cincinnati Genomics, Epigenomics and Sequencing Core and the University of Minnesota Genomics Center for their assistance in generating the methylation data for this project. Data for this study has been deposited into the Gene Expression Omnibus (GEO) under GSE197795 and will be released pending full publication of this manuscript.

## Supporting Information

**Supporting Tables:** S1-S5 Tables

**S1 Fig:**
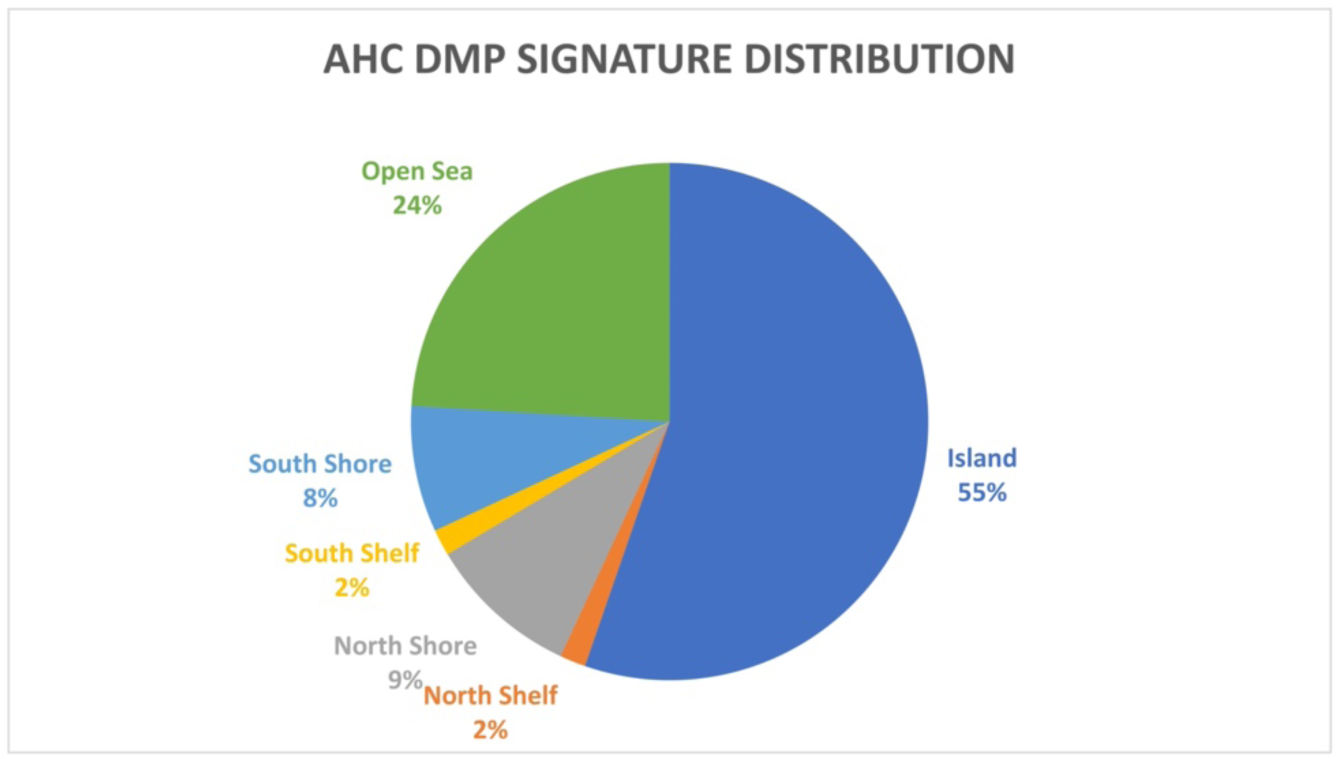
Genomic Distribution of DMPs within the AHC methylation profile. The distribution of 6,177 DMPs in the AHC profile. Most probes are found within CpG islands or closely associated with islands.

**S2 Fig:**
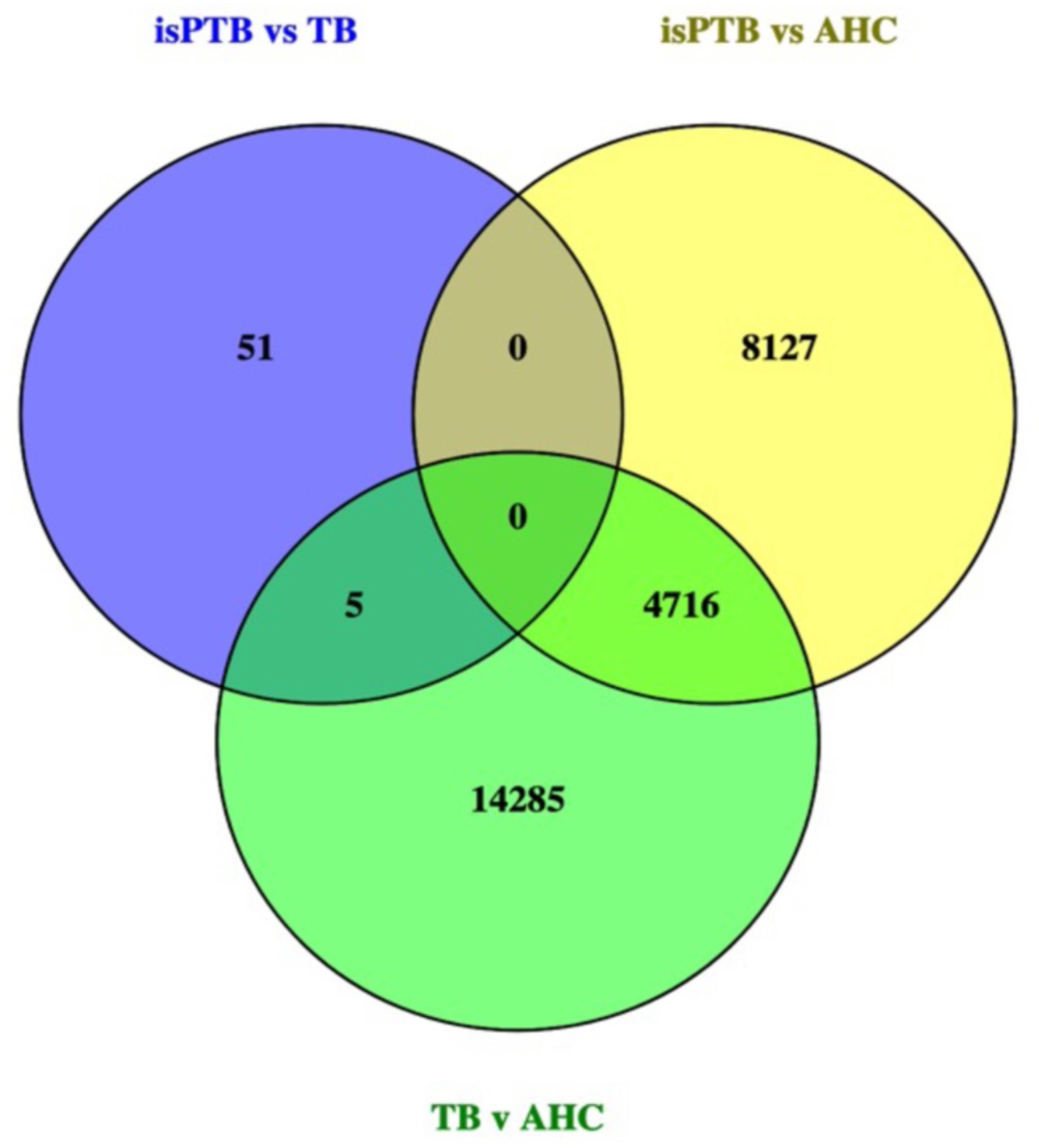
Intersection of significant DMRs. The venn diagram representing the intersection of pairwise comparisons to classify significant DMRs into isPTB and AHC specific profiles

## References

1. Blencowe H, Cousens S, Chou D, Oestergaard M, Say L, Moller A-B, et al. Born Too Soon: The global epidemiology of 15 million preterm births. Reproductive Health. 2013;10(Suppl 1):S2.

2. Chawanpaiboon S, Vogel JP, Moller A-B, Lumbiganon P, Petzold M, Hogan D, et al. Global, regional, and national estimates of levels of preterm birth in 2014: a systematic review and modelling analysis. The Lancet Global Health. 2018;7(Acta Obstet Gynecol Scand 56 1977):e37–46.

3. Monangi NK, Brockway HM, House M, Zhang G, Muglia LJ. The genetics of preterm birth: Progress and promise. Seminars in perinatology. 2015;39(8):574–83.

4. Burton GJ, Fowden AL. The placenta: a multifaceted, transient organ. Philosophical Transactions of the Royal Society of London B: Biological Sciences. 2015;370(1663):20140066.

5. Khong TY, Mooney EE, Ariel I, Balmus NCM, Boyd TK, Brundler M-A, et al. Sampling and Definitions of Placental Lesions: Amsterdam Placental Workshop Group Consensus Statement. Archives of Pathology & Laboratory Medicine. 2016;140(7):698–713.

6. Sankar DK, Bhanu SP, Kiran S, Ramakrishna B, Shanthi V. Vasculosyncytial membrane in relation to syncytial knots complicates the placenta in preeclampsia: a histomorphometrical study. Anatomy & Cell Biology. 2012;45(2):86–91.

7. Morgan TK. Role of the Placenta in Preterm Birth: A Review. American Journal of Perinatology. 2016;33(03):258–66.

8. Nijman TA, Vliet EO van, Benders MJ, Mol BW, Franx A, Nikkels PG, et al. Placental histology in spontaneous and indicated preterm birth: A case control study. Placenta. 2016;48:56–62.

9. Manuck TA, Esplin SM, Biggio J, Bukowski R, Parry S, Zhang H, et al. The phenotype of spontaneous preterm birth: application of a clinical phenotyping tool. American Journal of Obstetrics and Gynecology. 2015;212(4):487.e1-487.e11.

10. Benton SJ, Leavey K, Grynspan D, Cox BJ, Bainbridge SA. The clinical heterogeneity of preeclampsia is related to both placental gene expression and placental histopathology. American journal of obstetrics and gynecology. 2018;219(6):604.e1-604.e25.

11. Moutquin J-M. Classification and heterogeneity of preterm birth. BJOG: An International Journal of Obstetrics and Gynaecology. 2003;110(s20):30–3.

12. Yuen RKC, Robinson WP. Review: A high capacity of the human placenta for genetic and epigenetic variation: Implications for assessing pregnancy outcome. Placenta. 2011;32:S136–41.

13. Whigham C-AA, MacDonald TM, Walker SP, Hannan NJ, Tong S, Kaitu’u-Lino TJ. The untapped potential of placenta-enriched molecules for diagnostic and therapeutic development. Placenta. 2019;

14. Ghaemi MS, Tarca AL, Romero R, Stanley N, Fallahzadeh R, Tanada A, et al. Proteomic signatures predict preeclampsia in individual cohorts but not across cohorts – implications for clinical biomarker studies. J Maternal-fetal Neonatal Medicine. 2021;1–8.

15. Steyerberg EW. Clinical Prediction Models, A Practical Approach to Development, Validation, and Updating. Statistics Biology Heal. 2008;83–100.

16. Brockway HM, Kallapur SG, Buhimschi IA, Buhimschi CS, Ackerman WE, Muglia LJ, et al. Unique transcriptomic landscapes identified in idiopathic spontaneous and infection related preterm births compared to normal term births. PloS one. 2019;14(11):e0225062.

17. Yuen RK, Peñaherrera MS, Dadelszen P von, McFadden DE, Robinson WP. DNA methylation profiling of human placentas reveals promoter hypomethylation of multiple genes in early-onset preeclampsia. European journal of human genetics : EJHG. 2010;18(9):1006–12.

18. Parets SE, Conneely KN, Kilaru V, Menon R, Smith AK. DNA methylation provides insight into intergenerational risk for preterm birth in African Americans. Epigenetics. 2015;10(9):784–92.

19. Avila L, Yuen RK, Diego-Alvarez D, Peñaherrera MS, Jiang R, Robinson WP. Evaluating DNA methylation and gene expression variability in the human term placenta. Placenta. 2010;31(12):1070–7.

20. Wilson SL, Leavey K, Cox BJ, Robinson WP. Mining DNA methylation alterations towards a classification of placental pathologies. Human molecular genetics. 2018;27(1):135–46.

21. Gardiner-Garden M, Frommer M. CpG Islands in vertebrate genomes. J Mol Biol. 1987;196(2):261–82.

22. Maksimovic J, Phipson B, Oshlack A. A cross-package Bioconductor workflow for analysing methylation array data. F1000Research. 2016;5:1281.

23. Huber W, Carey VJ, Gentleman R, Anders S, Carlson M, Carvalho BS, et al. Orchestrating high-throughput genomic analysis with Bioconductor. Nat Methods. 2015;12(2):115–21.

24. Team Rs. RStudio: Integrated Development for R. RStudio. 2020; Available from: http://www.rstudio.com/.

25. Team RC. R: A language and environment for statistical ## computing. R Foundation for Statistical Computing [Internet]. 2020; Available from: https://www.R-project.org/

26. Aryee MJ, Jaffe AE, Corrada-Bravo H, Ladd-Acosta C, Feinberg AP, Hansen KD, et al. Minfi: a flexible and comprehensive Bioconductor package for the analysis of Infinium DNA methylation microarrays. Bioinformatics. 2014;30(10):1363–9.

27. S D, P D, S B, Triche, T Jr, M B. methylumi: Handle Illumina methylation data. 2020; Available from: https://www.bioconductor.org/packages/release/bioc/html/methylumi.html

28. Fortin J-P, Labbe A, Lemire M, Zanke BW, Hudson TJ, Fertig EJ, et al. Functional normalization of 450k methylation array data improves replication in large cancer studies. Genome Biology. 2014;15(11):503.

29. Zhou W, Laird PW, Shen H. Comprehensive characterization, annotation and innovative use of Infinium DNA methylation BeadChip probes. Nucleic Acids Research. 2017;45(4):e22–e22.

30. McCartney DL, Walker RM, Morris SW, McIntosh AM, Porteous DJ, Evans KL. Identification of polymorphic and off-target probe binding sites on the Illumina Infinium MethylationEPIC BeadChip. Genomics Data. 2016;9:22–4.

31. Amemiya HM, Kundaje A, Boyle AP. The ENCODE Blacklist: Identification of Problematic Regions of the Genome. Scientific Reports. 2019;9(1):9354.

32. Price EM, Robinson WP. Adjusting for Batch Effects in DNA Methylation Microarray Data, a Lesson Learned. Frontiers in genetics. 2018;9:83.

33. Du P, Zhang X, Huang C-C, Jafari N, Kibbe WA, Hou L, et al. Comparison of Beta-value and M-value methods for quantifying methylation levels by microarray analysis. BMC Bioinformatics. 2010;11(1):587.

34. Ritchie ME, Phipson B, Wu D, Hu Y, Law CW, Shi W, et al. limma powers differential expression analyses for RNA-sequencing and microarray studies. Nucleic Acids Research. 2015;43(7):e47–e47.

35. Benjamini Y, Hochberg Y. Controlling the False Discovery Rate: A Practical and Powerful Approach to Multiple Testing. Journal of the Royal Statistical Society: Series B (Methodological). 1995;57(1):289–300.

36. Oliveros, J.C. Venny. An interactive tool for comparing lists with Venn’s diagrams. 2015; Available from: https://bioinfogp.cnb.csic.es/tools/venny/index.html

37. Peters TJ, Buckley MJ, Statham AL, Pidsley R, Samaras K, Lord RV, et al. De novo identification of differentially methylated regions in the human genome. Epigenetics & Chromatin. 2015;8(1):6.

38. Mi H, Huang X, Muruganujan A, Tang H, Mills C, Kang D, et al. PANTHER version 11: expanded annotation data from Gene Ontology and Reactome pathways, and data analysis tool enhancements. Nucleic Acids Research. 2017;45(D1):D183–9.

39. Li D, Xie Z, Pape M, Dye T. An evaluation of statistical methods for DNA methylation microarray data analysis. BMC Bioinformatics. 2015;16(1):217.

40. Mansell G, Gorrie-Stone TJ, Bao Y, Kumari M, Schalkwyk LS, Mill J, et al. Guidance for DNA methylation studies: statistical insights from the Illumina EPIC array. Bmc Genomics. 2019;20(1):366.

41. Monteagudo-Sánchez A, Sánchez-Delgado M, Mora JRH, Santamaría NT, Gratacós E, Esteller M, et al. Differences in expression rather than methylation at placenta-specific imprinted loci is associated with intrauterine growth restriction. Clin Epigenetics. 2019;11(1):35.

42. Wilson SL, Robinson WP. Utility of DNA methylation to assess placental health. Placenta. 2018;64 Suppl 1:S23–8.

43. Wilson AS, Power BE, Molloy PL. DNA hypomethylation and human diseases. Biochimica Et Biophysica Acta Bba - Rev Cancer. 2007;1775(1):138–62.

44. Ehrlich M. DNA hypermethylation in disease: mechanisms and clinical relevance. Epigenetics. 2019;14(12):1–23.

45. Filion GJP, Zhenilo S, Salozhin S, Yamada D, Prokhortchouk E, Defossez P-A. A Family of Human Zinc Finger Proteins That Bind Methylated DNA and Repress Transcription. Mol Cell Biol. 2006;26(1):169–81.

46. Yu Y, Shang R, Chen Y, Li J, Liang Z, Hu J, et al. Tumor suppressive ZBTB4 inhibits cell growth by regulating cell cycle progression and apoptosis in Ewing sarcoma. Biomed Pharmacother. 2018;100:108–15.

47. Weber A, Marquardt J, Elzi D, Forster N, Starke S, Glaum A, et al. Zbtb4 represses transcription of P21CIP1 and controls the cellular response to p53 activation. Embo J. 2008;27(11):1563–74.

48. Paquette AG, Brockway HM, Price ND, Muglia LJ. Comparative transcriptomic analysis of human placentae at term and preterm delivery. Biology of reproduction. 2018;98(1):89–101.

49. Hirota Y, Daikoku T, Tranguch S, Xie H, Bradshaw HB, Dey SK. Uterine-specific p53 deficiency confers premature uterine senescence and promotes preterm birth in mice. Journal of Clinical Investigation. 2010;120(3):803–15.

50. Scales SJ, Hesser BA, Masuda ES, Scheller RH. Amisyn, a Novel Syntaxin-binding Protein That May Regulate SNARE Complex Assembly*. J Biol Chem. 2002;277(31):28271–9.

51. Wang T, Li L, Hong W. SNARE proteins in membrane trafficking. Traffic. 2017;18(12):767–75.

52. Han J, Pluhackova K, Böckmann RA. The Multifaceted Role of SNARE Proteins in Membrane Fusion. Front Physiol. 2017;8:5.

53. Bogaart G van den, Jahn R. Counting the SNAREs needed for membrane fusion. J Mol Cell Biol. 2011;3(4):204–5.

54. Guček A, Gandasi NR, Omar-Hmeadi M, Bakke M, Døskeland SO, Tengholm A, et al. Fusion pore regulation by cAMP/Epac2 controls cargo release during insulin exocytosis. Elife. 2019;8:e41711.

55. Rentero C, Puigdomènech P. Specific use of start codons and cellular localization of splice variants of human phosphodiesterase 9A gene. Bmc Mol Biol. 2006;7(1):39.

56. Gerbaud P, Taskén K, Pidoux G. Spatiotemporal regulation of cAMP signaling controls the human trophoblast fusion. Front Pharmacol. 2015;6:202.

57. Lim JH, Kim SY, Park SY, Lee SY, Kim MJ, Han YJ, et al. Non-Invasive Epigenetic Detection of Fetal Trisomy 21 in First Trimester Maternal Plasma. Plos One. 2011;6(11):e27709.

58. Frendo JL, Vidaud M, Guibourdenche J, Luton D, Muller F, Bellet D, et al. Defect of villous cytotrophoblast differentiation into syncytiotrophoblast in Down’s syndrome. The Journal of clinical endocrinology and metabolism. 2000;85(10):3700–7.

59. Pidoux G, Gerbaud P, Cocquebert M, Segond N, Badet J, Fournier T, et al. Review: Human trophoblast fusion and differentiation: Lessons from trisomy 21 placenta. Placenta. 2011;33:S81–6.

60. Malassiné A, Frendo J-L, Evain-Brion D. Trisomy 21- affected placentas highlight prerequisite factors for human trophoblast fusion and differentiation. International Journal of Developmental Biology. 2009;54(2– 3):475–82.

61. Bryant AH, Bevan RJ, Spencer-Harty S, Scott LM, Jones RH, Thornton CA. Expression and function of NOD-like receptors by human term gestation-associated tissues. Placenta. 2017;58:25–32.

62. Lappas M. NOD1 and NOD2 Regulate Proinflammatory and Prolabor Mediators in Human Fetal Membranes and Myometrium via Nuclear Factor-Kappa B. Biol Reprod. 2013;89(1):Article 14, 1-11.

63. Bourhis LL, Benko S, Girardin SE. Nod1 and Nod2 in innate immunity and human inflammatory disorders. Biochem Soc T. 2007;35(6):1479–84.

64. Marchand M, Horcajadas JA, Esteban FJ, McElroy SL, Fisher SJ, Giudice LC. Transcriptomic Signature of Trophoblast Differentiation in a Human Embryonic Stem Cell Model. Biology of Reproduction. 2011;84(6):1258–71.

65. Wang B, Palomares K, Parobchak N, Cece J, Rosen M, Nguyen A, et al. Glucocorticoid Receptor Signaling Contributes to Constitutive Activation of the Noncanonical NF-κB Pathway in Term Human Placenta. Molecular Endocrinology. 2013;27(2):203–11.

66. Tang D, Kang R, Coyne CB, Zeh HJ, Lotze MT. PAMPs and DAMPs: signal 0s that spur autophagy and immunity. Immunological Reviews. 2012;249(1):158–75.

67. Rijn BB van, Franx A, Steegers EAP, Groot CJM de, Bertina RM, Pasterkamp G, et al. Maternal TLR4 and NOD2 Gene Variants, Pro-Inflammatory Phenotype and Susceptibility to Early-Onset Preeclampsia and HELLP Syndrome. Plos One. 2008;3(4):e1865.

68. Robertson SA, Hutchinson MR, Rice KC, Chin P, Moldenhauer LM, Stark MJ, et al. Targeting Toll-like receptor-4 to tackle preterm birth and fetal inflammatory injury. Clin Transl Immunol. 2020;9(4):e1121.

69. Costello MJ, Joyce SK, Abrahams VM. NOD Protein Expression and Function in First Trimester Trophoblast Cells. Am J Reprod Immunol. 2007;57(1):67–80.

70. Strauss JF, Romero R, Gomez-Lopez N, Haymond-Thornburg H, Modi BP, Teves ME, et al. Spontaneous preterm birth: advances toward the discovery of genetic predisposition. Am J Obstet Gynecol. 2018;218(3):294–314.e2.

71. Härtel Ch, Finas D, Ahrens P, Kattner E, Schaible Th, Müller D, et al. Polymorphisms of genes involved in innate immunity: association with preterm delivery. Mhr Basic Sci Reproductive Medicine. 2004;10(12):911–5.

72. Tekola-Ayele F, Zeng X, Ouidir M, Workalemahu T, Zhang C, Delahaye F, et al. DNA methylation loci in placenta associated with birthweight and expression of genes relevant for early development and adult diseases. Clin Epigenetics. 2020;12(1):78.

73. Vennou KE, Kontou PI, Braliou GG, Bagos PG. Meta-analysis of gene expression profiles in preeclampsia. Pregnancy Hypertens. 2020;19:52–60.

74. Ramasammy R, Munisammy L, Sweta K, Selvakumar S, Velu K, Rani J, et al. Association between GCK gene polymorphism and gestational diabetes mellitus and its pregnancy outcomes. Meta Gene. 2021;28:100856.

75. Beaumont RN, Warrington NM, Cavadino A, Tyrrell J, Nodzenski M, Horikoshi M, et al. Genome wide associatin study of offspring birth weight in 86577 women identifies five novel loci and highlights maternal genetic effects that are independent of fetal genetics. Human Molecular Genetics. 2018;27(4):742–56.

76. Lang CT, Markham KB, Behrendt NJ, Suarez AA, Samuels P, Vandre DD, et al. Placental Dysferlin Expression is Reduced in Severe Preeclampsia. Placenta. 2009;30(8):711–8.

77. Varanou A, Withington SL, Lakasing L, Williamson C, Burton GJ, Hemberger M. The importance of cysteine cathepsin proteases for placental development. J Mol Med. 2006;84(4):305–17.

78. Jia R-Z, Zhang X, Hu P, Liu X-M, Hua X-D, Wang X, et al. Screening for differential methylation status in human placenta in preeclampsia using a CpG island plus promoter microarray. Int J Mol Med. 2012;30(1):133– 41.

79. Uhlén M, Fagerberg L, Hallström BM, Lindskog C, Oksvold P, Mardinoglu A, et al. Tissue-based map of the human proteome. Science. 2015;347(6220):1260419.

80. Guo X, Zhang Y, Wang P, Li T, Fu W, Mo X, et al. VSTM1-v2, a novel soluble glycoprotein, promotes the differentiation and activation of Th17 cells. Cell Immunol. 2012;278(1–2):136–42.

81. Sonderegger S, Pollheimer J, Knöfler M. Wnt Signalling in Implantation, Decidualisation and Placental Differentiation – Review. Placenta. 2010;31(10):839–47.

82. Knöfler M, Pollheimer J. Human placental trophoblast invasion and differentiation: a particular focus on Wnt signaling. Frontiers in Genetics. 2013;4:190.

83. Kokkinos MI, Murthi P, Wafai R, Thompson EW, Newgreen DF. Cadherins in the human placenta – epithelial–mesenchymal transition (EMT) and placental development. Placenta. 2010;31(9):747–55.

84. Adu-Gyamfi EA, Czika A, Gorleku PN, Ullah A, Panhwar Z, Ruan L-L, et al. The Involvement of Cell Adhesion Molecules, Tight Junctions, and Gap Junctions in Human Placentation. Reprod Sci. 2021;28(2):305– 20.

85. Yeung KR, Chiu CL, Pidsley R, Makris A, Hennessy A, Lind JM. DNA methylation profiles in preeclampsia and healthy control placentas. Am J Physiol-heart C. 2016;310(10):H1295–303.

86. Gal H, Lysenko M, Stroganov S, Vadai E, Youssef SA, Tzadikevitch-Geffen K, et al. Molecular pathways of senescence regulate placental structure and function. Embo J. 2019;38(18):e100849.

87. He S, Gu W, Li Y, Zhu H. ANRIL/CDKN2B-AS shows two-stage clade-specific evolution and becomes conserved after transposon insertions in simians. Bmc Evol Biol. 2013;13(1):247.

88. Gamage T, Schierding W, Hurley D, Tsai P, Ludgate JL, Bhoothpur C, et al. The role of DNA methylation in human trophoblast differentiation. Epigenetics. 2018;13(12):1154–73.

89. Mayne BT, Leemaqz SY, Smith AK, Breen J, Roberts CT, Bianco-Miotto T. Accelerated placental aging in early onset preeclampsia pregnancies identified by DNA methylation. Epigenomics-uk. 2016;9(3):279–89.

90. Raj K, Horvath S. Current perspectives on the cellular and molecular features of epigenetic ageing. Exp Biol Med. 2020;245(17):1532–42.

91. Lee Y, Choufani S, Weksberg R, Wilson SL, Yuan V, Burt A, et al. Placental epigenetic clocks: estimating gestational age using placental DNA methylation levels. Aging. 2019;11(12):4238–53.

