## Supplemental Fiiles for "Characterization of methylation profiles in spontaneous preterm birth placental villous tissue"

**S1 Table: Probe Filtering during quality control assessment**

| **Total probes read into pipeline** | | **866,901** |
|---|---|---|
| Failed Detection | 810 | 866,091 |
| Normalization (FunNorm) | 232 | 865,859 |
| **Filtering probes after quality control** | | |
| Probes that failed in 2+ samples | 8,546 | **865,859** |
| Remove X/Y probes | 18,913 | 857,313 |
| Remove SNPs (Manifest) | 28,488 | 838,400 |
| Remove SNPs (Zhou 2016) | 13,303 | 809,912 |
| Remove Cross hybridizing probes (McCartney 2016) | 38,280 | 796,609 |
| Remove Blacklist probes (2019 Blacklist) | 119 | 758,329 |
| **Total probes left for analyses** | | **758,210** |

**S2 Table: Statistical testing in limma to determine significant DMPs between pairwise comparisons**

| | **BH adjusted p <0.05** | | | **BH adjusted p <0.1** | | |
|---|---|---|---|---|---|---|
| | **isPTB vs AHC** | **TB vs AHC** | **isPTB vs TB** | **isPTB vs AHC** | **TB vs AHC** | **isPTB vs TB** |
| **More methylated probes** | 13,111 | 41,767 | 0 | 27,797 | 71,566 | 0 |
| **Less methylated probes** | 17,037 | 31,632 | 0 | 36,791 | 59,535 | 0 |
| **Total DMPs** | 30,148 | 73,399 | 0 | 64,588 | 131,101 | 0 |

| | **BH adjusted p <0.2** | | | **BH adjusted p <0.3** | | |
|---|---|---|---|---|---|---|
| | **isPTB vs AHC** | **TB vs AHC** | **isPTB vs TB** | **isPTB vs AHC** | **TB vs AHC** | **isPTB vs TB** |
| **More methylated probes** | 51,382 | 116,150 | 0 | 73,338 | 152,935 | 29 |
| **Less methylated probes** | 72,136 | 109,987 | 7 | 105,617 | 152,647 | 593 |
| **Total DMPs** | 123,518 | 226,137 | 7 | 178,955 | 305,582 | 662 |

*”Separate” was selected as the statistical method within limma

** Limma only selected for adjusted p-value, not log2 fold-change.

**S3 Table: Statistical testing in DMRcate to determine significant DMPs between pairwise comparisons**

| BH adjusted p | <0.05 | <0.2 | <0.3 | <0.5 |
|---|---|---|---|---|
| **isPTB vs TB** | 0 | 7 | 662 | 14,611 |
| **isPTB vs AHC** | 30,148 | 123,518 | 178,955 | 300,625 |
| **TB vs AHC** | 73,399 | 226,137 | 305,582 | 483,450 |

** Limma only selected for adjusted p-value, not log2 fold-change.

**S4 Table: The top 25 differentially DMR mean differences in pairwise comparisons**

| DMR location | Locus Name | Mean Diff AHC vs TB | Mean Diff AHC vs isPTB | Mean Diff TB vs isPTB |
|---|---|---|---|---|
| chr2:11915711-11916260 | ***MIR3681HG*** | 0.1474 | 0.1065 | Not significant |
| chr1:150692971-150694343 | ***GOLPH3L*** | 0.1366 | 0.0874 | Not significant |
| chr22:19973978-19975691 | ***ARVCF*** | 0.1125 | 0.0887 | Not significant |
| chr16:85342729-85343936 | ***GSE1*** | 0.1023 | 0.0615 | Not significant |
| chr9:34372089-34373067 | ***MYORG*** | 0.0993 | 0.0698 | Not significant |
| chr19:6230050-6230665 | ***MLLT1*** | 0.0986 | 0.0519 | Not significant |
| chr4:12224743-12225077 | ***LINC02270*** | 0.0946 | 0.0679 | Not significant |
| chr8:103750821-103751623 | ***RIMS2*** | 0.0894 | 0.0832 | Not significant |
| chr2:794646-796536 | ***LINC01115*** | 0.0881 | 0.0683 | Not significant |
| chr10:121577971-121579007 | ***FGFR2*** | 0.0812 | 0.0774 | Not significant |
| chr13:45965025-45966279 | ***ZC3H13*** | 0.0800 | 0.0565 | Not significant |
| chr22:46440394-46442103 | ***CELSR1*** | 0.0786 | 0.0848 | Not significant |
| chr19:54040774-54041856 | ***VSTM1*** | 0.0753 | 0.0491 | Not significant |
| chr11:62211493-62212431 | ***SCGB2A1*** | 0.0748 | 0.0606 | Not significant |
| chr4:7967275-7969643 | ***ABLIM2*** | 0.0741 | 0.0827 | Not significant |
| chr12:75057893-75058468 | ***KCNC2*** | 0.0723 | 0.0780 | Not significant |
| chr12:126018024-126018364 | ***AC005186.1*** | 0.0721 | 0.0576 | Not significant |
| chr16:89488412-89489377 | ***ANKRD11*** | 0.0714 | 0.0426 | Not significant |
| chr19:13616871-13617970 | ***CACNA1A*** | 0.0678 | 0.0609 | Not significant |
| chr1:41831580-41832649 | ***HIVEP3*** | 0.0668 | 0.0545 | Not significant |

| | | | | |
|---|---|---|---|---|
| chr7:65878352-65879115 | ***VKORC1L1*** | 0.0667 | 0.0491 | Not significant |
| chr17:66097276-66098113 | ***CEP112*** | 0.0665 | 0.0687 | Not significant |
| chr3:195619562-195620147 | ***MUC20P1*** | 0.0653 | 0.0550 | Not significant |
| chr2:43327937-43328914 | ***THADA*** | 0.0647 | 0.0588 | Not significant |
| chr7:44152238-44154322 | ***GCK*** | 0.0632 | 0.0475 | Not significant |
| chr17:46018654-46019184 | ***MAPT*** | -0.0470 | -0.0518 | Not significant |
| chr19:44302666-44303858 | ***ZNF235*** | -0.0494 | -0.0474 | Not significant |
| chr6:161560605-161561121 | ***PRKN*** | -0.0494 | -0.0327 | Not significant |
| chr16:67184164-67185527 | ***EXOC3L1*** | -0.0506 | -0.0433 | Not significant |
| chr11:17568197-17569556 | ***OTOG*** | -0.0522 | -0.0294 | Not significant |
| chr22:23744094-23745131 | ***ZNF70*** | -0.0536 | -0.0400 | Not significant |
| chr1:160084263-160085568 | ***KCNJ9*** | -0.0541 | -0.0222 | Not significant |
| chr8:123859056-123859953 | ***FER1L6*** | -0.0563 | -0.0473 | Not significant |
| chr16:8724073-8724983 | ***ABAT*** | -0.0577 | -0.0662 | Not significant |
| chr15:78933106-78934580 | ***CTSH*** | -0.0579 | -0.0794 | Not significant |
| chr11:129993525-129993935 | ***PRDM10*** | -0.0596 | -0.0476 | Not significant |
| chr17:27312855-27313499 | ***WSB1*** | -0.0598 | -0.0373 | Not significant |
| chr19:30413468-30414886 | ***ZNF536*** | -0.0621 | -0.0370 | Not significant |
| chr20:25013229-25014771 | ***ACSS1*** | -0.0649 | -0.0353 | Not significant |
| chr16:30485296-30485966 | ***ITGAL*** | -0.0683 | -0.0504 | Not significant |
| chr1:1296671-1297807 | ***ACAP3*** | -0.0685 | -0.0662 | Not significant |
| chr2:11679584-11680144 | ***LPIN1*** | -0.0691 | -0.0437 | Not significant |
| chr19:14048977-14049823 | ***IL27RA*** | -0.0702 | -0.0460 | Not significant |
| chr9:123656764-123657427 | ***DENND1A*** | -0.0794 | -0.0852 | Not significant |
| chr16:31366142-31366536 | ***ITGAX*** | -0.0852 | -0.0428 | Not significant |
| chr7:133811022-133812369 | ***EXOC4*** | -0.0945 | -0.0578 | Not significant |
| chr12:69724920-69725444 | ***AC025263.1*** | -0.0988 | -0.0738 | Not significant |
| chr15:90208739-90209326 | ***SEMA4B*** | -0.1083 | -0.1185 | Not significant |
| chr22:24988020-24990749 | ***KIAA1671*** | -0.1093 | -0.0773 | Not significant |
| chr10:93334974-93335677 | ***MYOF*** | -0.1173 | -0.0643 | Not significant |

**S5 Table: Comparison of Methylation and Transcription profiles for intersected AHC DMRs**

| | | | Methylation profile | | | Transcription Profile | | |
|---|---|---|---|---|---|---|---|---|
| **DMR Genomic Location** | **DMR Associated Gene** | **Total CpGs in DMR** | **AHC vs TB*** | **AHC vs isPTB*** | **PTB vs TB*** | **AHC vs TB Log2 Fold Change **** | **AHC vs isPTB Log2 Fold Change**** | **isPTB vs TB Log2 Fold Change **** |
| chr9:21993972-21995735 | ***CDKN2A-CDKN2B-AS*** | 13 | -0.0126 | -0.0116 | #N/S | 1.17 | 0.88 | 0.29 |
| chr12:6938111-6939048 | ***ATN1*** | 6 | -0.0177 | -0.0081 | #N/S | 1.12 | 0.93 | 0.19 |
| chr22:41939981-41941494 | ***CENPM*** | 14 | -0.0126 | -0.0058 | #N/S | 1.24 | 1.4 | -0.15 |
| chr7:108095719-108097606 | ***LAMB4*** | 11 | -0.0360 | -0.0167 | #N/S | -1.08 | -0.96 | -0.12 |
| chr16:23680392-23681287 | ***PLK1*** | 5 | 0.0296 | 0.0327 | #N/S | 1.15 | 1.11 | 0.04 |
| chr15:40731625-40735036 | ***RAD51*** | 15 | -0.0192 | -0.0142 | #N/S | 1.07 | 1.22 | -0.14 |
| chr15:64752519-64753130 | ***RBPMS2*** | 6 | -0.0159 | -0.0155 | #N/S | 1.57 | 1.17 | 0.39 |
| chr22:24180492-24181665 | ***SUSD2*** | 11 | -0.0124 | -0.0053 | #N/S | 1.41 | 1.62 | -0.21 |

***Mean Difference in Methylation over all CpGs in the DMR**
****Differential Expression of Gene as reported in(16)  #N/S = Not significant**
